# Axonal TAU sorting requires the C-terminus of TAU but is independent of ANKG and TRIM46 enrichment at the AIS

**DOI:** 10.1101/2020.06.26.173526

**Authors:** M. Bell, S. Bachmann, J. Klimek, F. Langerscheidt, H. Zempel

## Abstract

Somatodendritic missorting of the axonal protein TAU is a hallmark of Alzheimer’s disease and related tauopathies. Cultured rodent primary neurons and iPSC-derived neurons are used for studying mechanisms of neuronal polarity, including TAU trafficking. However, these models are expensive, time-consuming and/or require the sacrification of animals. In this study, we evaluated four differentiation procedures to generate mature neuron cultures from human SH-SY5Y neuroblastoma cells, in comparison to mouse primary neurons, and tested their TAU sorting capacity. We show that SH-SY5Y-derived neurons, differentiated with sequential RA/BDNF treatment, are suitable for investigating axonal TAU sorting. These human neurons show pronounced neuronal polarity, axodendritic outgrowth, expression of the neuronal maturation markers TAU and MAP2, and, importantly, efficient axonal sorting of endogenous and transfected human wild type TAU, similar to primary neurons. We demonstrate that axonal TAU enrichment requires the presence of the C-terminal half, as a C-terminus-lacking construct (N-term-TAU^HA^) is not axonally enriched in both neuronal cell models. Moreover, SH-SY5Y-derived neurons do not show formation of a classical axon initial segment (AIS), indicated by the lack of Ankyrin G (ANKG) and tripartite motif-containing protein 46 (TRIM46) at the proximal axon, which suggests that successful axonal TAU sorting is independent of classical AIS formation. Taken together, our results suggest i) that SH-SY5Y-derived neurons are a valuable human neuronal cell model for studying TAU sorting, which is readily accessible at low cost and without animal need, and that ii) the mechanisms of axonal TAU targeting require the TAU C-terminal half but are independent of ANKG or TRIM46 enrichment at the proximal axon.

## Introduction

The microtubule-associated protein TAU is highly abundant in the axons of healthy human brain neurons^1,2^, where six isoforms are predominantly expressed^3,4^. By binding microtubule (MT) filaments, TAU regulates their assembly and disassembly^5–7^, and is involved in axonal outgrowth, plasticity, MT-dependent cargo transport and other cellular functions^8–12^. The C-terminal repeat domains and their flanking regions bind the MTs^13,14^, while the N-terminus acts as a spacer between adjacent MT filaments and mediates membrane interactions^15–17^. Missorting of highly phosphorylated TAU into the somatodendritic compartment and the formation of insoluble TAU aggregates (‘neurofibrillary tangles’) are major hallmarks of Alzheimer’s disease (AD) and related tauopathies^18–23^. Subcellular TAU mislocalization leads to a lack of axonal TAU-MT interaction, depletion of stable MT filaments and, eventually, to disruption of axonal transport^24–26^. In the dendrites, abnormal amounts of hyperphosphorylated TAU trigger TTLL6-mediated MT destabilization and spine loss, FYN-mediated excitotoxicity, and other processes leading to postsynaptic degradation^27–32^.

Understanding the mechanisms of axonal TAU sorting in healthy neurons is a critical step towards the elucidation of pathological processes that accompany TAU missorting. Several mechanisms are thought to drive axonal TAU sorting^33,34^, including active anterograde TAU trafficking, selective axonal TAU-MT interactions, or active retrograde retention of axonal TAU. In this context, the axon initial segment (AIS), a key player for developing and maintaining neuronal polarity with Ankyrin G (ANKG) as a master organizer of the molecular structure^35–37^, may be relevant, as polarized MT filaments at the AIS are involved in motor protein-mediated soma-to-axon-trafficking^35,38^, presumably including TAU transit^33,34^. The tripartite motif containing-protein 46 (TRIM46) may also be involved into anterograde TAU sorting by regulating the MT architecture at the AIS^39^. Moreover, recent studies found direct evidence for a retrograde TAU diffusion barrier within the AIS that is compromised by MT disassembly^40,41^. Notably, little is yet known about the intrinsic components of TAU, i.e. functional domains, interaction motifs, posttranslational modifications and relevant TAU interactions, although some recent studies in rodent primary neurons assumed major roles for phosphorylation sites within the Proline-rich region and for the very N-terminal domain of TAU, but only a minor role for the four MT-binding repeat domains^42,43^.

Which neuronal cell models are available to address these key aspects of TAU sorting and missorting? Two commonly used cellular systems are rodent primary neuron cultures^40–43^ and recently also human induced pluripotent stem cell (iPSC)-derived neurons^44–48^. Both systems have considerable limitations. Rodent primary neuronal cultures require the use and sacrification of animals, and rodent TAU differs in several aspects from human TAU, e.g. regarding the isoform expression pattern^49^. In addition, the human cellular environment, i.e. the entire sorting machinery and interaction partners may differ from that in humans. Overexpressed human TAU often appears uniformly distributed in rodent primary neurons^50^. IPSC-derived neurons overcome many of these issues, but their generation is expensive, time-consuming and results in mixed cultures with variable homogeneity and differentiation efficiency^51–53^. The neuroblastoma cell line SH-SY5Y is human-derived^54^ and cells can be differentiated with various substances into polarized neuronal cells (SH-SY5Y-derived neurons) at low cost^55^. SH-SY5Y-derived neurons express common neuronal maturation marker proteins^56–61^, including all six major human TAU isoforms^62,63^, show proper axonal TAU localization, and mimic the phosphorylation state of TAU in the adult human brain^64–67^. In brief, SH-SY5Y-derived neurons unite the benefits of a cell line – i.e. robust maintenance, fast growth rates, the saving of animal resources and simple accessibility for stable genetic engineering^68,69^ – with the higher scientific relevance of primary and iPSC derived neurons. In this study, we evaluated four differentiation procedures of SH-SY5Y cells for generating neuronal cells and tested their suitability for the investigation of TAU sorting. Mouse primary forebrain neurons were used as a reference neuronal cell model. We show that RA/BDNF-treated SH-SY5Y-derived neurons achieve pronounced cellular polarity, axonal outgrowth, strong upregulation of the neuronal markers TAU and MT-associated protein 2 (MAP2), and, importantly, efficient axonal sorting of endogenous TAU and overexpressed human TAU (0N3R-TAU^HA^). In contrast, TAU without the C-terminal half (N-term-TAU^HA^) is not axonally enriched in SH-SY5Y-derived and mouse primary neurons. Of note, SH-SY5Y-derived neurons do not show enrichment of ANKG and TRIM46 at the proximal axon despite successful axonal TAU targeting, suggesting an ANKG- and TRIM46-independent mechanism of axonal TAU enrichment. In brief, our results give evidence that i) SH-SY5Y-derived neurons show successful axonal sorting of endogenous and transfected human TAU, ii) the N-terminus of TAU is not sufficient for proper axonal sorting, and ii) the axonal targeting of endogenous and 0N3R-TAU^HA^ is independent of ANKG, a master organizer of the AIS molecular structure, and TRIM46, a AIS-located MT-organizing protein.

## Methods

### Molecular biology

For experiments with endogenous and transfected TAU, an expression vector, which harbored the cDNA encoding for the tdTomato fluorescent protein, was used as volume marker control. For experiments with recombinant TAU constructs, N-terminally tagged 0N3R-TAU^HA^ and N-term-TAU^HA^ constructs were cloned into the same, CMV promoter-driven expression vector (see Suppl. Sequence 1). For this, the intended TAU cDNA sequences were amplified by PCR using primer pairs with restriction sites (AgeI, BamHI) corresponding to the multiple cloning site of the backbone vector. Additionally, the forward primer contained a human influenza hemagglutinin (HA) tag sequence overhang with start codon (5’-ATGTACCCATACGATGTTCCAGATTACGCT-3’), which was fused to the 5’ site of the TAU sequence. The PCR product was analyzed via gel electrophoresis, and correctly sized inserts were cut out and purified with NucleoSpin™ Gel & PCR Clean up Kit (Macherey & Nagel). Vector and insert were digested (30 min, 37 °C) according to manufacturer’s protocols (enzymes by New England Biolabs), either purified directly (insert) or from agarose gel (vector) and ligated (16-18 h, 20-22 °C) with T4 DNA ligase (NEB) according to the manufacturer’s protocol.

For plasmid amplification, plasmids were transformed into chemocompetent bacteria (*E. coli* Top10™, Thermo Fisher Scientific, TFS), which were generated like previously described^70^. Bacteria were spread onto pre-warmed agar plates with antibiotics (16-18 h, 37 °C), colonies were picked and grown in nutrient medium with antibiotics. For plasmid isolation and purification from bacterial suspensions, the PureYield™ MidiPrep Kit (ProMega) and NucleoBond Xtra™ Midi/Maxi Kit (Macherey & Nagel) were used according to the manufacturer’s protocols. Plasmid concentration and purity were checked spectrophotometrically (NanoDrop™ 100, TFS) and isolates were long-term stored at −20 °C.

### SH-SY5Y cell cultivation and differentiation

#### Maintenance

The SH-SY5Y neuroblastoma cell line was kindly provided by Prof. Dr. Rudolf Wiesner (Institute of Veg. Physiology II, University Hospital Cologne). SH-SY5Y cells were cultured in DMEM/F12 (#10565-018, TFS) supplemented with 10 % fetal bovine serum (FBS, Biochrom AG), penicillin/streptomycin and amphotericin B (1X Anti/Anti, #15240062, TFS) (referred to as ‘SHM-10’) on uncoated cell culture flasks (VWR). Cultures were maintained in a sterile incubator (MCO-20AIC, PHCbi) at 37 °C, 95 % air humidity and 5 % CO_2_ concentration. Culture medium was changed once or twice per week, depending on cell density, and cells were passaged for further cultivation or differentiation experiments at 70-80 % confluency. For passaging, SH-SY5Y cells were washed with DPBS, trypsinized (0.05 % Trypsin/EDTA, TFS) for 3-5 minutes, spun down at 1000 g for 3 minutes and resuspended in SHM-10. For differentiation experiments, resuspended SH-SY5Y cells were counted with an automatic cell counter (TC20™, Bio-Rad) and seeded with 7.5 to 10 x 10^3^ cells/cm^2^ onto glass coverslips (VWR), coated with 20 μg/ml Poly-D-lysine (PDL, AppliChem) for at least 3 h at 37 °C. For long-term storage, trypsinized and spun down cells were resuspended in FBS with 10 % DMSO, cooled down with −1 °C/min in a cryo container (Mr. Frosty™, VWR) and stored at −80 °C or in liquid nitrogen.

#### Differentiation

Differentiation protocols were started 24 h after seeding. For RA-based differentiation, fresh SMM with 10 μM retinoic acid (RA, Sigma-Aldrich) was added at day 0 (d0) and replaced every 2-3 days up to d7. Differentiation with RA & the brain-derived neurotrophic factor (BDNF, Peprotech) was adapted from former protocols^56^. Briefly, after RA treatment up to d7 as described above, cells were washed once with DPBS, and SHM-10 medium without FBS (referred to as ‘SHM-0’) containing 10 ng/ml BDNF was added and once replaced until d14. Differentiation protocols for Phorbol-12-myristate-13-acetate (TPA)- and RA/TPA-based differentiation were adapted from former protocols^60,71^. In brief, fresh SHM-10 with 81 μM TPA (Sigma-Aldrich) was added and replaced every 2-3 days until d9 for TPA-based differentiation. For RA & TPA-treatment, fresh SHM-10 with 10 μM RA was added, replaced after 2-3 days up to d4 or d5, whereupon cells were washed once with PBS and cultivated in fresh SHM-10 with 81 μM TPA until d9 with one medium change in between.

Two alternative long-term differentiation procedures were tested for RA, RA/TPA and TPA treatments. The first variant (variant A) included cultivation with RA (for RA & RA/TPA protocols) and TPA (TPA protocol) in SHM-10 until d7, cultivation with RA (RA protocol) and TPA (RA/TPA & TPA protocols) in SHM-10 with 3 % FBS (SHM-3) until d9, and cultivation with RA (RA protocol) and TPA (RA/TPA & TPA protocols) in SHM-0 until d14. The second variant (variant B), where the seeding density was increased to 15 to 20 x 10^3^ cells/cm^2^, included cultivation with RA (for RA & RA/TPA protocols) and TPA (TPA protocol) in SHM-10 until d2, cultivation with RA (RA & RA/TPA protocols) and TPA (TPA protocol) in SHM-3 until d7, cultivation with RA (RA protocol) and TPA (RA/TPA & TPA protocols) in SHM-3 until d9, and cultivation with RA (RA protocol) and TPA (RA/TPA & TPA protocols) in SHM-0 until d14. The acquired cultures showed decreased viability after transfection (see Suppl.Fig. 1).

### Transfection

SH-SY5Y-derived neurons were transfected using the polymer-based PolyJet™ DNA transfection reagent (#SL100688, SignaGen) at d5 of differentiation. Transfection was performed according to the manufacturer’s protocol with the following modifications: Before transfection, conditioned medium was collected from each well and stored at 37 °C with 5 % CO_2_. For one well of a 24-well plate, 0.33 μg of plasmid DNA and 1 μl PolyJet™ reagent were separately prepared with 25 μl DMEM (#P04-03500, TFS), then mixed, incubated and added dropwise to the cultures. Medium was changed to previously collected, conditioned medium, supplemented with fresh doses of RA or TPA, 3-16 h after transfection. Transfection efficiency ranged from 5-10 %. For experiments with recombinant TAU, tdTomato- and TAU^HA^ construct-encoding plasmids were co-transfected in SH-SY5Y-derived neurons. For this, DNA of both plasmids was incubated commonly with PolyJet™. The plasmid DNA was mixed with a ratio of three (TAU^HA^ plasmid) to one part (tdTomato plasmid). The total DNA amount per well was not altered, i.e. 0.33 μg was used for one well of a 24-well plate.

### Primary neurons cultivation

#### Isolation

Pregnant female FVB wild type mice were anesthetized and sacrificed at day 13.5 of pregnancy by authorized personnel with respect to the official regularities and guidelines of the governmental authority, the Landesumweltamt (LANUV) of North Rhine-Westphalia, Germany. Preparation of E13.5 embryos, and isolation and cultivation of primary neurons were performed like previously described^72^. In brief, embryos were decapitated, embryonic scalp, skull and meninges were removed, the cerebral hemispheres were isolated, washed, dissociated for 5-7 minutes with 0.05 % Trypsin/EDTA, and homogenized in HBSS (w/o Ca^2+^/Mg^2+^, #14185052, TFS). Dissociated primary cells were counted as described above for SH-SY5Y cells and seeded with 5 to 7.5 x 10^4^ cells/cm^2^ in neuronal plating medium (NPM), i.e. Neurobasal™ medium (#21103, TFS) supplemented with 1 % FBS, penicillin/streptomycin and amphotericin B (1X Anti/Anti), GlutaMAX™ (1X, TFS) and NS-21 (1X, #P07-20001, PAN Biotech), onto glass coverslips, coated with 20 μg/ml PDL for at least 3 h at 37 °C.

#### Maintenance

Four days after seeding, the medium was doubled by adding neuronal maintenance medium (NMM), i.e. NPM without FBS. Cytosine arabinoside (Ara-C, #C1768, Sigma-Aldrich) was added to an end concentration of 0.5 – 1.0 μg/ml to impair survival of glial, endothelial and other proliferating cells.

#### Transfection

Mouse primary forebrain neurons were transfected at day 6 in vitro (div6). Transfection was performed as described previously^73^. In brief, 0.25 μg of plasmid DNA and 0.375 μl Lipofectamine™ 2000 (#743517, Invitrogen) were mixed for one well of a 24-well plate, incubated for 30 minutes and added dropwise to the cultures. Medium was changed to previously collected, conditioned medium 60 minutes after transfection. For experiments with recombinant TAU, tdTomato- and TAU^HA^ construct-encoding plasmids were co-transfected in mouse primary neurons. For this, the DNA of both plasmids was mixed with a ratio of three parts (TAU^HA^ DNA) to one part (tdTomato DNA) before addition of Lipofectamine™ 2000. The total DNA amount per well was not altered, i.e. 0.25 μg per 24-well (1 μg per 6-well) was used for one well of a 24-well plate.

### Fixation & Immunofluorescence

Fixation and immunofluorescence of cell cultures was done as described previously^73^ with slight modifications. In brief, cells were fixed for 1 h with 3.7 % formaldehyde (in PBS), permeabilized and blocked with 5 % bovine serum albumin (BSA, Sigma-Aldrich) and 0.1 % Triton X-100 (AppliChem) in PBS for 5 minutes, incubated with the primary antibody (diluted in PBS) either for 2-3 h at 20-22 °C or preferentially at 4 °C for 16-18 h, thoroughly washed, and incubated with the corresponding secondary antibody (diluted in PBS), which was coupled to AlexaFluor™ fluorophores, for 1-2 h at 20-22 °C. For detection of ANKG and TRIM46 levels, the fixation process was modified (30 minutes instead of 1 h) according to the sensitivity of AIS proteins to fixation procedures reported in previous studies^74,75^. The following primary antibodies were used: rabbit anti-TAU (K9JA) (1:1000, #A0024, DAKO), rabbit anti-HA (1:1000, #3724S, cellSignaling), mouse anti-HA (1:1000, #901533, BioLegend), chicken anti-MAP2 (1:2000, ab5392, Abcam), mouse anti-ANKG (N106/36) (1:250, MABN466, Neuromab), rabbit anti-TRIM46 (1:500, #377003, Synaptic Systems). Nuclei were stained with NucBlue™ (1 drop/ml, Hoechst 33342, TFS) for 20-30 minutes, and samples were mounted on objective slides (#145-0011, Bio-Rad) using aqueous PolyMount™ (#18606, Polysciences) or Mowiol™ (AppliChem) mounting medium, dried for 24 h at RT and long-term stored at 4 °C in the dark.

### Microscopy

Immunostained cells were imaged with a widefield fluorescence microscope (Axioscope 5, Zeiss), using an LED excitation lamp (Colibri 7, Zeiss), a fluorescence camera device (Axiocam 503 mono, Zeiss), objectives with 10 x, 20 x, 40 x (air-based) and 63 x (oil-based) magnification (Zeiss) and Zen imaging software (Zen Blue pro, Zeiss). Exposure time and light intensity were adjusted to avoid excessive sample bleaching and to preclude saturation of fluorescent signal detection. For the analysis of protein expression levels, all images were taken with identical settings regarding laser intensity and exposure time to ensure statistical comparability. For documentation during cell cultivation, images were taken with a brightfield light microscope (DM IL LED, Leica) using objectives with 10x and 20 x (air-based) magnification (Leica) and Las X imaging software (Leica).

### Data analysis

#### Differentiation Efficiency

The efficiency of SH-SY5Y cell differentiation was defined as relative number of SH-SY5Y-derived neurons compared to undifferentiated and non-neuronal cells. SH-SY5Y cells were considered as differentiated if they fulfilled the following morphological criteria: i) roundish shaped soma, ii) changed nuclear morphology, i.e. smaller and brighter nuclear signal, iii) axonal outgrowth exceeds soma diameter at least twofold and iv) determined axon length exceeds 50 μm. Further analysis was performed only with cells reaching these criteria.

#### Axonal outgrowth

Axonal outgrowth of SH-SY5Y-derived neurons was determined using the Fiji/ImageJ software (NIH). The processes were traced manually, and axon length was determined as pixel number, transformed to absolute distances with respect to the original image resolution. The measured section reached from the soma-to-axon-border to the most distal axonal part, which was still detectable. Included cells fulfilled the described differentiation criteria and the axon was distinctive from processes of adjacent cells.

#### Endogenous protein expression

For the analysis of total endogenous expression of the neuronal maturation markers TAU and MAP2, SH-SY5Y-derived neurons were analyzed with Fiji/ImageJ software. Entire cells were encircled manually, and the mean fluorescence intensity (MFI_neuron_) of this marked region of interest (ROI) was determined. To exclude background fluorescence, MFI of an empty area next to the neuron was determined in the same manner (MFI_noise_) and subtracted from the MFI_neuron_. Further, MFI was determined in absence of primary antibodies (MFI_unspecific_), and also subtracted from MFI_neuron_ to eliminate impact of unspecific binding of the secondary antibody. Protein levels of differentiated cells were normalized to that of undifferentiated cells. To ensure comparability of protein levels, all cell cultures of one experiment were cultivated on the same plate, immunostained and mounted in parallel with identical antibody solutions and imaged with identical microscope settings.

#### Axonal enrichment of endogenous TAU

By using Fiji/ImageJ, a random patch within the soma, which had no overlap with the nuclear signal, was set as first ROI for each analyzed SH-SY5Y-derived neuron. An axonal segment of around 5 – 10 μm in 50 – 75 μm distance to the soma was chosen as second ROI. TAU signals were measured as MFI for both somatic and axonal ROIs (MFI_soma_, MFI_axon_). To exclude background fluorescence, MFI of empty areas next to the soma and axon were determined in the same manner (MFI_noiseSoma_, MFI_noiseAxon_) and subtracted from the MFI_soma_/MFI_axon_. The ratio of MFI_axon_ and MFI_soma_ was calculated (axon-to-soma-ratio). The axon-to-soma-ratio of the volume marker tdTomato was determined using identical ROIs and identical calculation steps. The axon-to-soma-ratio of TAU was normalized to this of tdTomato to achieve the axonal enrichment factor of TAU (AEF_TAU_). Only tdTomato-positive SH-SY5Y-derived neurons were included for analysis.

#### Axonal enrichment of TAU^HA^ constructs

ROI selection and calculation of the axonal enrichment factor of TAU^HA^ constructs (AEF_TAU_HA_) were done as described above for endogenous TAU with a few deviations. Instead of TAU signals, the intensity of anti-HA signal was measured. Only cells that were successfully co-transfected with tdTomato and TAU^HA^ were included for analysis. Further, to overcome signal of unspecific anti-HA antibody binding, MFIs of somatic and axonal ROIs from untransfected cells (MFI_unspecificSoma_, MFI_unspecificAxon_) were measured and averaged for each experiment and subtracted from all MFIs of TAU^HA^ from transfected cells.

#### Plot profiles of the proximal axon

The expression profiles of ANKG, TRIM46 and MAP2 proteins were determined at the proximal axon of SH-SY5Y-derived and mouse primary neurons. By using the Fiji/ImageJ software, the first 75 μm of the axon, starting with the soma-to-axon-border, were traced with a segmented line (thickness: three pixels), and a plot profile was generated. To exclude influence of background fluorescence, a plot profile of the region right next to the axon was measured and subtracted from the original value. Further, a plot profile was determined in absence of primary antibodies and subtracted from the sample values to eliminate impact of unspecific binding of the secondary antibody.

#### Protein enrichment at the proximal axon

For the calculation of the protein enrichment at the proximal axon, the section of the mean plot profiles, where the fluorescent signal reached at least 50 % of the maximum value, was determined in mouse primary neurons. All values within this section were averaged, separately for each individual experiment. In order to achieve the enrichment factor of ANKG andTRIM46 at the proximal axon, this average value was normalized against a baseline value of each experiment. This baseline value was acquired by averaging the most distal values of the plot profile. Here, the number of included values was equal to the value number of the previously determined section at the proximal axon. The identical sections used for mouse primary neurons were then applied to the corresponding plot profiles of SH-SY5Y-derived neurons to determine the protein enrichment of ANKG and TRIM46 in the same manner.

#### Statistical analysis

All statistical calculations were performed using PRISM™ analysis software (V8.3, GraphPad Inc.). For comparisons between one group of samples and a constant number (e.g. 1.0) or between two groups of samples, a two-sided student’s t-test was used for the determination of p-values. If the standard deviation differed significantly among the two groups (p ≥ 0.05), the t-test was conducted with Welch’s correction. When multiple groups were tested for significant differences, a Brown-Forsythe-test was applied to check if the standard deviation (SD) differed significantly (p ≥ 0.05) among the tested groups. If this was the case, a Brown-Forsythe and Welch’s ANOVA test with Dunnett’s T3 correction for multiple comparisons was conducted. If the SD was not significantly different (p < 0.05), an ordinary one-way ANOVA test with Tukey’s correction for multiple comparisons was conducted. The chosen statistical tests are mentioned for all experiments in the corresponding figure caption. The Gaussian distribution of all data sets was confirmed by a Shapiro-Wilk normality test prior to statistical analyses. Significance levels are symbolized with asterisks if p < 0.05 (*), p < 0.01 (**) or p < 0.001 (***), p < 0.0001 (****).

## Results

### RA/BDNF treatment produces homogeneous cultures of highly polarized neurons

We first aimed to investigate whether SH-SY5Y-derived neurons are suitable for studying TAU sorting. For this, naïve human SH-SY5Y neuroblastoma cells (Fig. 1A) were differentiated using four distinct procedures (Fig. 1B). Cells were either treated only with retinoic acid (RA) until day 7 of differentiation (d7), with RA until d7 followed by brain-derived neurotrophic factor (BDNF) in serum-free conditions until d14, with the phorbol ester TPA until d9 or with RA until d4 followed by TPA until d9. To compare the amount of successfully differentiated neurons, cells were classified as differentiated when they fulfilled several criteria, e.g. roundish shaped somata and axonal processes of at least 50 μm length (see methods for details). While treatment with RA (52.5 % ± 11.8 % neurons), TPA (39.7 % ± 10.2 %), and RA/TPA (47.4 % ± 5.4 %) generated mixed cultures of non-neuronal and neuronal cells, the sequential RA/BDNF treatment resulted in high numbers of SH-SY5Y-derived neurons (74.1 % ± 11.4 %) (Fig. 1C&D).

**Figure 1:**
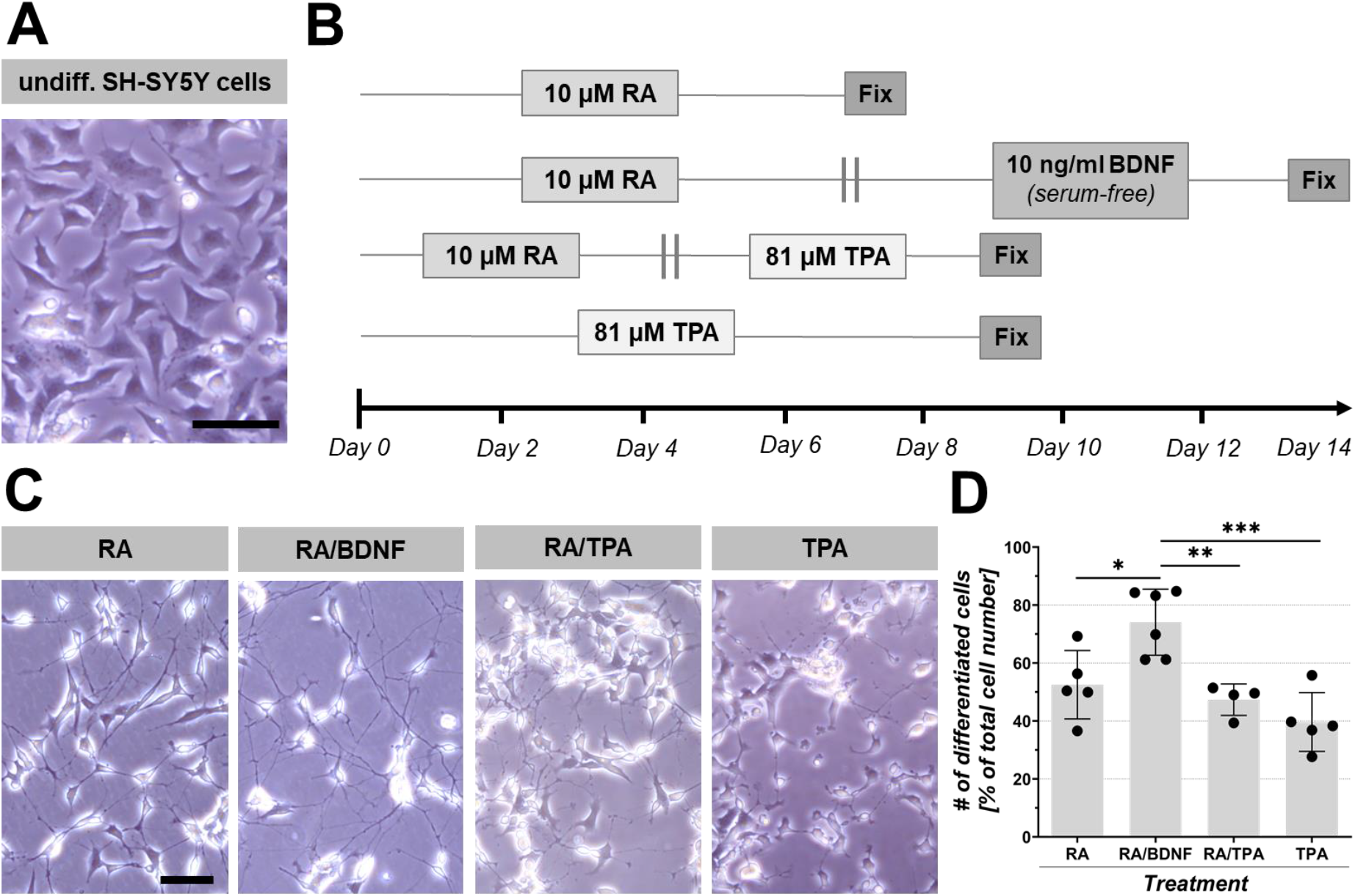
Generation of SH-SY5Y-derived neurons. Representative images of undifferentiated SH-SY5Y cells, timeline of differentiation procedures (see methods for details), and differentiated SH-SY5Y-derived neurons. **A:** Undifferentiated SH-SY5Y cells cultured in normal SHM-10 medium. Scale bar: 50 μm. **B:** Schematic time course of differentiation protocols. The bottom line indicates the day (d) of differentiation. **B.1:** RA was applied until d7 in SH-SY5Y cell culture medium with 10 % FBS (SHM-10). **B.2:** Sequential treatment with RA in SHM-10 until d7 and BDNF in serum-free SHM-0 medium until d14. **B.3:** Sequential treatment of RA in SHM-10 until d4 and TPA in SHM-10 until d9. **B.4:** Application of TPA in SHM-10 until d9. Grey doubled lines indicate substance changes, grey box represents fixation of cells. **C:** SH-SY5Y-derived neurons after differentiation with RA, RA/BDNF, RA/TPA or TPA in culture. Note i) the neurite outgrowth of SH-SY5Y-derived neurons upon all treatments and ii) the different extent of co-cultured undifferentiated cells. Scale bar: 50 μm. **E:** Quantification of differentiation efficiency. Quantification was done for four to six independent experiments with 1000 ± 50 cells per experiment. Differentiated cells were determined with the differentiation criteria (see methods for details). Black dots represent independent experiments, grey bars indicate the arithmetic mean of all experiments, and error bars show SD. An ordinary one-way ANOVA with Tukey’s correction for multiple comparisons was performed to determine significance levels between all four methods. Significance levels: * p < 0.05, ** p < 0.01, *** p < 0.001.

### RA/BDNF-treated SH-SY5Y-derived neurons show pronounced axonal outgrowth and high levels of neuronal maturation markers

Next, the neuronal maturity of SH-SY5Y-derived neurons upon the four treatments was evaluated morphologically and biochemically. The axonal outgrowth of SH-SY5Y-derived neurons was determined as the distance from the soma-to-axon-border to the distal axonal end. The average axonal outgrowth in RA/BDNF-induced neurons (264 μm ± 43 μm) exceeded the axon length of all other cultures significantly, namely RA (141 μm ± 21 μm), RA/TPA (125 μm ± 28 μm), and TPA (76 μm ± 10 μm), indicating the highest level of neuronal polarity in RA/BDNF-induced SH-SY5Y-derived neurons (Fig. 2A&C). Next, we tested the expression levels of two commonly used marker proteins for neuronal maturation^76^, TAU and the somatodendritic MT-associated protein 2 (MAP2) (Fig. 2B). The expression of TAU was upregulated 12-fold (± 4.6) in RA/BDNF-treated neurons compared to undifferentiated SH-SY5Y cells. A similar increase of TAU protein levels was observed upon RA/TPA treatment (9.6-fold ± 2.8). Treatment only with RA (6.4-fold ± 3.3) or TPA (5.6-fold ± 2.6) led to less strong increases of TAU levels (Fig. 2D). MAP2 levels were strongly elevated in neurons after RA/BDNF treatment (8.8-fold ± 3.6), with notable inter-experimental variability, and moderately elevated after differentiation with RA (3.7-fold ± 1.6) and RA/TPA (4.1-fold ± 2.2). Treatment with TPA (1.8-fold ± 1.1) did not result in neurons with enhanced MAP2 levels (Fig. 2E). The attempt to cultivate SH-SY5Y cells in RA, TPA or RA/TPA under serum-reduced conditions were not successful, as cultures largely died upon the transfection procedure (Suppl.Fig. 1C-H), in contrast to transfected RA/BDNF-treated cultures (Suppl.Fig. 1A&B).

**Fig. 2:**
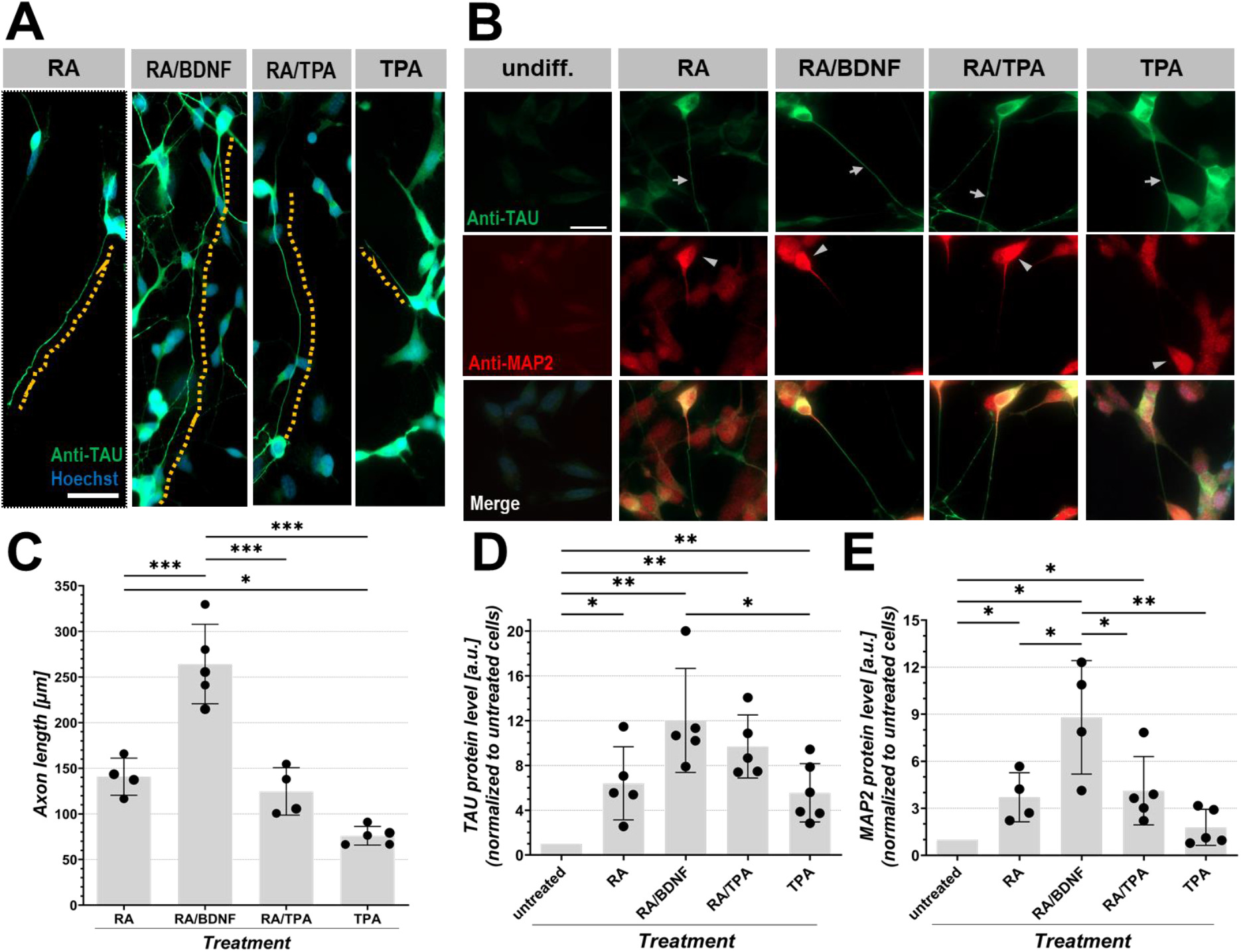
Neuronal maturation of SH-SY5Y-derived neurons upon differentiation. **A:** Axonal outgrowth of SH-SY5Y-derived neurons after differentiation with RA, RA/BDNF, RA/TPA or TPA, immunostained with polyclonal anti-TAU (K9JA) antibody and NucBlue™ (nuclear staining, Hoechst 33342). Dashed lines trace representative axons. Note the different extent of axonal elongation. Scale bar: 50 μm. **B:** Representative images of undifferentiated SH-SY5Y and SH-SY5Y-derived neurons immunostained with polyclonal anti-TAU (K9JA) and polyclonal anti-MAP2 antibodies. TAU (green) and MAP2 (red) are shown in single channels and merged. Note the weak signal of neuronal maturation markers TAU (green) and MAP2 (red) in undifferentiated cells (left panel) and the enhanced signals after differentiation as indicated. Scale bar: 20 μm; **C-E:** Analysis of neuronal maturation in SH-SY5Y-derived neurons, which were determined by nuclear and somatic shape, and neuritic outgrowth (see differentiation criteria (methods) for details). Quantification of axonal outgrowth (C) was done for four to five independent experiments with 30-35 cells traced per experiment. Quantification for TAU (D) and MAP2 (E) protein levels was done for five to six (D) or four to five (E) independent experiments with 25-30 cells per experiment. Black dots represent independent experiments, grey bars indicate the arithmetic mean of all experiments, and error bars show SD. An ordinary one-way ANOVA with Tukey’s correction for multiple comparisons was performed to determine significance levels between all four treatments (C), and all four treatments and undifferentiated cells (D&E). Significance levels: * p < 0.05, ** p < 0.01, *** p < 0.001.

Taken together, SH-SY5Y-derived neurons, which were treated with RA/BDNF, showed the highest differentiation efficiency and pronounced neuronal maturation, indicated by axonal outgrowth and the highest expression levels of TAU and MAP2 protein.

### Endogenous TAU is efficiently sorted in SH-SY5Y-derived neurons upon RA/BDNF treatment

To further evaluate the suitability of SH-SY5Y-derived neurons for TAU sorting studies, the efficiency of endogenous TAU sorting was determined. For this, the cells were transfected with tdTomato-expressing plasmids at day 5 of differentiation (d5), and the axonal enrichment factor of TAU (AEF_endoTAU_) was defined as axon-to-soma-ratio of TAU, normalized to the axon-to-soma-ratio of the randomly distributing volume marker tdTomato (see methods for details).

RA/BDNF-treated neurons show mainly axonal TAU and strong somatic tdTomato signals (Fig. 3A), indicating efficient axonal sorting of TAU (AEF_endoTAU_ = 7.9 ± 1.2) (Fig. 3B). For all other treatments, the SH-SY5Y-derived neurons show less axonal TAU sorting, which results in more yellowish axons in the merged images (Fig. 3A), and lower AEF_endoTAU_ values for RA/TPA- (3.3 ± 0.6), TPA- (3.4 ± 0.8) and RA-treated neurons (Fig. 3B). The AEF_endoTAU_ value of RA/BDNF-treated SH-SY5Y-derived neurons hints at the efficient intracellular sorting of TAU protein.

**Figure 3:**
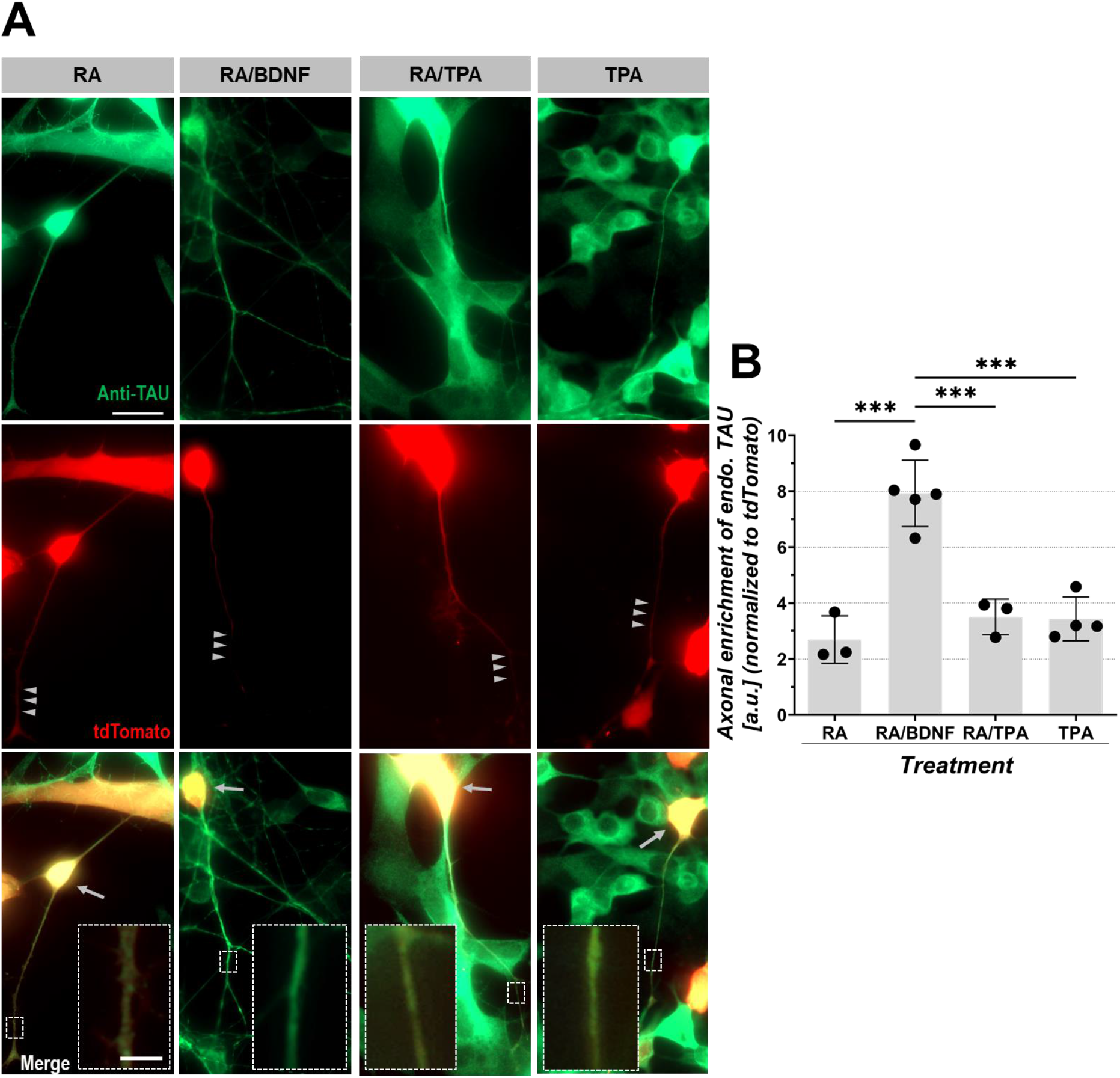
Axonal enrichment of endogenous TAU in SH-SY5Y-derived neurons. SH-SY5Y-derived neurons were transfected with tdTomato and immunostained with a polyclonal anti-TAU (K9JA) antibody. **A:** Endogenous TAU (green) and transfected tdTomato as volume marker (red) are shown in single channels and merged. Note the strong axonal sorting of TAU compared to tdTomato in SH-SY5Y-derived neurons after RA/BDNF treatment (arrowheads), which leads to yellowish somatic staining (arrows) and mainly green axonal TAU staining in merged images. Axonal sections (small dashed boxes) are shown with higher magnification (large dashed boxes) in merged images. Scale bar: 20 μm, scale bar within dashed box: 3 μm. **B:** Quantification of the axonal enrichment of endogenous TAU (AEF_endoTAU_), normalized to intracellular tdTomato distribution in neuronal cells (see methods for details). Quantification was done for three to five independent experiments with 20-30 cells per experiment. Black dots represent independent experiments, grey bars indicate the arithmetic mean of all experiments, and error bars show SD. An ordinary one-way ANOVA with Tukey’s correction for multiple comparisons was performed to determine significance levels between all conditions. Significance level: *** p < 0.001.

Since the results of the comparative analyses revealed RA/BDNF-treated SH-SY5Y-derived neurons as the most suitable system in many respects, i.e. we observed high numbers of neurons, pronounced neuronal polarity, efficient sorting of endogenous TAU, and unchanged viability up to nine days after transfection, only those neurons were used for all further experiments.

### SH-SY5Y-derived neurons show robust sorting of overexpressed 0N3R-TAU^HA^

Efficient axonal sorting of transfected and overexpressed TAU constructs is an often-faced problem in primary neuron models^50^. Therefore, we checked whether SH-SY5Y-derived neurons achieve an endogenous-like sorting efficiency of transfected human wild type TAU. For this, we co-transfected SH-SY5Y-derived neurons with tdTomato and the shortest TAU isoform 0N3R, fused to an N-terminal HA tag (0N3R-TAU^HA^) (Fig. 4B), at d5 of differentiation. The subcellular distribution was again quantified at d14 (Fig. 4A&C). The same experiments were done in mouse primary forebrain neurons, with co-transfection at div6 and quantification at div9 (Fig. 4D).

**Figure 4:**
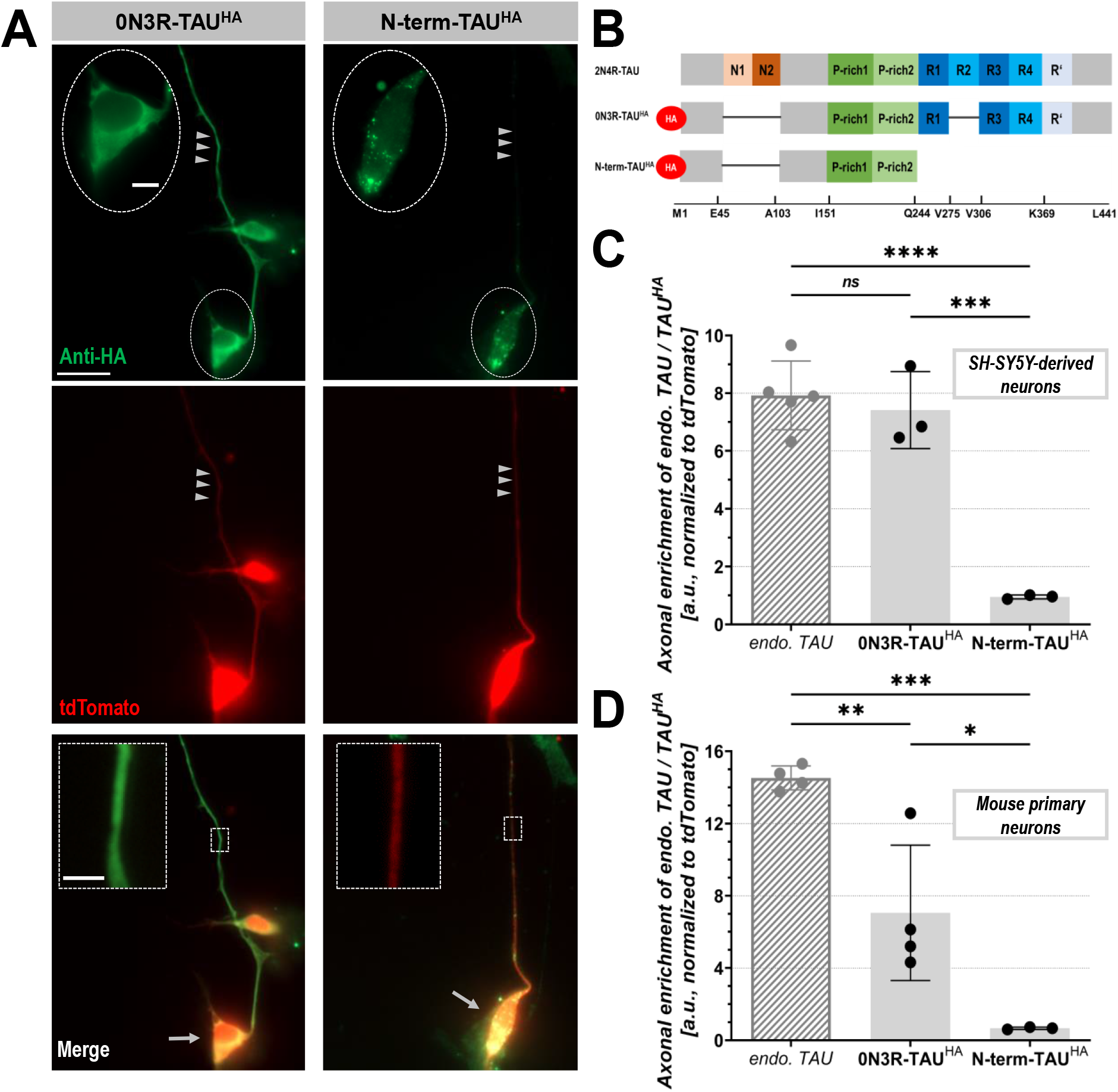
Axonal enrichment of transfected TAU^HA^ constructs in RA/BDNF-treated SH-SY5Y-derived and mouse primary neurons. **A:** Representative immunofluorescent images of SH-SY5Y-derived neurons transfected with tdTomato and either 0N3R-TAU^HA^ (left column) or N-term-TAU^HA^ (right column). TAU^HA^ constructs (green) and co-transfected tdTomato as volume marker (red) are shown in single channels and merged. Cells were transfected at d5 and fixed at d14 (see methods for details). Note the strong axonal sorting of TAU compared to tdTomato (arrowheads) in 0N3R-TAU^HA^ which leads to yellowish somatic staining (arrows) and mainly green axonal staining in merged images. Axons of cells with N-term-TAU^HA^ appear red, indicating weak axonal sorting of N-term-TAU^HA^. Axonal sections (small dashed boxes) are shown with higher magnification (large dashed boxes). HA-positive puncta (small dashed circle, magnified in large dashed circle) appear in N-term-TAU^HA^ transfected cells but not in cells expressing 0N3R-TAU^HA^. Scale bar: 20 μm, scale bar within dashed circle: 5 μm, scale bar within dashed box: 3 μm. **B:** Domain structure of full-length 2N4R-TAU compared to the used TAU constructs. 0N3R-TAU^HA^ contains the amino acids M1 – E45, A103 – V275 and V306 – L441. N-term-TAU^HA^ contains the amino acids M1 – E45 and A103 – V244. Missing parts are shown as black lines. Both constructs were N-terminally fused to an HA tag (red circle). The bottom line indicates selected amino acid positions. N1/N2: N-terminal inserts, P-rich domain: Proline-rich domain, R1 – R4: C-terminal repeat domains. R’: Pseudo-repeat domain. **C&D:** Quantification of axonal enrichment of TAU^HA^ constructs (AEF_TAU_HA_) in SH-SY5Y-derived neurons (C) and mouse primary neurons (D, see Fig. S2 for corresponding images), normalized to intracellular tdTomato distribution. Quantification was done for three independent experiments with 20-55 cells per experiment (C), or for three to four independent experiments with 16-46 cells per experiment (D). The quantification of AEF_endoTAU_ in SH-SY5Y-derived neurons is included for comparison (striped bar, see Fig. 3A). Black dots represent independent experiments, grey bars indicate the arithmetic mean of all experiments, and error bars show SD. An ordinary one-way ANOVA with Tukey’s correction for multiple comparisons was performed to determine significance levels between endogenous TAU and the TAU^HA^ constructs. Significance levels: * p < 0.05, ** p < 0.01, *** p < 0.001, **** p < 0.0001, ns: p ≥ 0.05.

Strikingly, the axonal enrichment of the overexpressed 0N3R-TAU^HA^ construct (AEF_TAU_HA_ = 7.4 ± 1.3) was similar to that of endogenous TAU in RA/BDNF-treated SH-SY5Y-derived neurons at d14 (~ 94 % of AEF_endoTAU_ = 7.9 ± 1.2) (Fig. 4A&C). In mouse primary neurons, 0N3R-TAU^HA^ was also axonally sorted (AEF_TAU_HA_ = 7.1 ± 3.2), but less efficient than the endogenous mouse TAU (~ 49 % of AEF_endoTAU_ = 14.5 ± 0.6) (Fig. 4D, Suppl.Fig. 2). These results demonstrate that SH-SY5Y-derived neurons are able to sort overexpressed TAU with similar efficiency as endogenous TAU. This makes SH-SY5Y-derived neurons suitable for addressing key questions of TAU sorting behavior, such as the role of TAU domains, modifications, or interactions within the proximal axon.

### TAU^HA^ lacking the C-terminus shows no axonal sorting in SH-SY5Y-derived and mouse primary neurons

In order to demonstrate this suitability, we next evaluated the sorting behavior of a truncated TAU^HA^ version. As former data suggested an important role of the N-terminus for axonal sorting^43^, we used a construct which lacks the entire C-terminal half (N-term-TAU^HA^) (Fig. 4B). The sorting efficiency of N-term-TAU^HA^ was analyzed in SH-SY5Y-derived and mouse primary neurons. Interestingly, N-term-TAU^HA^ was axonally enriched neither in SH-SY5Y-derived neurons (AEF_TAU_HA_ = 0.97 ± 0.03) nor in mouse primary neurons (AEF_TAU_HA_ = 0.67 ± 0.05) (Fig. 4A,C,D, Suppl.Fig. 2). Moreover, N-term-TAU^HA^ tended to accumulate in the soma, indicated by HA-positive puncta in some neurons (Fig. 4A, Suppl.Fig. 2). These results, which are consistent in both used neuronal model systems, strongly suggest that the N-terminal half of TAU is either not involved in the process of axonal sorting or at least not sufficient for its successful implementation.

### ANKG and TRIM46 are not enriched at the AIS region in SH-SY5Y-derived neurons despite efficient axonal TAU sorting

Our data illustrate the pronounced neuronal polarity of SH-SY5Y-derived neurons, including the efficient axonal targeting of endogenous and transfected TAU. Since the axon initial segment (AIS) is essential for the development and maintenance of neuronal polarity^35,37^, we evaluated the enrichment of two AIS proteins, Ankyrin G (ANKG) and tripartite motif-containing protein 46 (TRIM46), in these cells (Fig. 5). ANKG is a master organizer of the AIS architecture as it recruits AIS-located scaffold proteins, ion channels or cytoskeletal elements^35,37,38^. TRIM46 is at MT-organizing protein that was shown to enrich at the proximal axon prior to ANKG and to promote the polarized distribution of TAU^39,77^.

**Figure 5:**
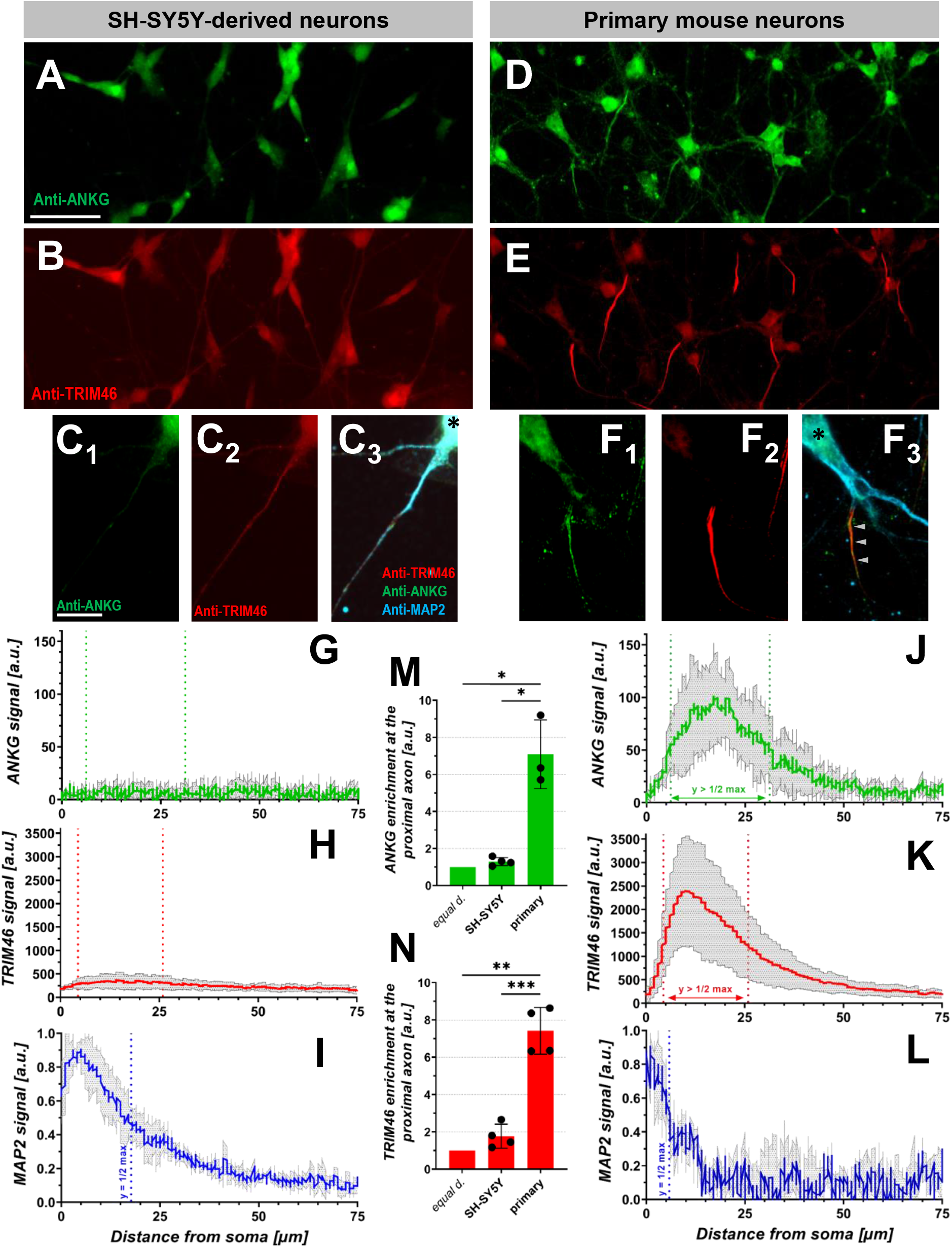
Formation of the AIS in SH-SY5Y-derived and mouse primary neurons. **A-F:** Representative images of ANKG and TRIM46 expression in SH-SY5Y-derived neurons (A-C) and mouse primary neurons (D-F), immunostained with monoclonal anti-ANKG (N106/36), polyclonal rabbit anti-TRIM46, and polyclonal anti-MAP2 antibodies. **A,B&D,E:** Overview of immunostained neuronal cultures. Scale bar: 50 μm. **C&F:** Magnification of one immunostained SH-SY5Y-derived (C) and mouse primary neuron (F). Note the clearly defined enrichment of TRIM46 and ANKG at the proximal axon of mouse primary neurons (arrowheads, F3), which is not visible at the proximal axon (axon defined by decreasing MAP2 intensity and process outgrowth >75 μm) of SH-SY5Y-derived neurons. Asterisks indicate nucleus position. Scale bar (A): 20 μm, scale bar (C.1): 5 μm. **G-L:** Plot profiles of TRIM46 (G&J), ANKG (H&K), and MAP2 (I&L) signals at the proximal axon of SH-SY5Y-derived (G-I) and mouse primary neurons (J-L). The colored lines indicate the arithmetic mean of three to four independent experiments with 28-36 traced cells per experiment (see methods for details). The grey area shows SD. **G,H&J,K:** The green and red dotted lines mark the corridor where the ANKG/TRIM46 signal are above 50 % of the maximum value (determined for the signals of mouse primary neurons in J&K). **I&L:** The MAP2 plot profiles were normalized to 1 prior to analysis. The blue dotted lines indicate the distance where the MAP2 signal reaches 50 % of the maximum value. **M&N:** Enrichment of ANKG (M) and TRIM46 (N) at the proximal axon (i.e. within the marked corridor), normalized to the baseline (see methods for details). Equal d. = equal distribution (reference value = 1). Quantification was done for three to four independent experiments with 28-36 cells per experiment. Black dots represent independent experiments, colored bars indicate the arithmetic mean of all experiments, and error bars show SD. A two-sided t-test was performed to determine significance levels between the cell models and 1. A two-sided t-test with Welch’s correction was performed to determine significance levels between both cell models. Significance levels: * p < 0.05, ** p < 0.01, *** p < 0.001.

In mouse primary neurons, we observed robust expression of TRIM46 and ANKG at the proximal axon, already at div9 (Fig. 5D-K) with a strong enrichment of both proteins at the proximal axon (Fig. 5M&N). In addition to TAU (Fig. 4D, Suppl. Fig. 2), also MAP2 shows a polarized subcellular distribution, as it is tightly restricted to the somatodendritic compartment (Fig. 5L). This is in line with previous studies on neuronal polarity that suggest the ANKG-dependent AIS formation to be necessary for somatic MAP2 restriction^35,39,78^. The somatic retention of MAP2 is less efficient in SH-SY5Y-derived neurons, indicated by the more than 3-fold increased distance of the half-maximal intensity (SH-SY5Y: 17.7 μm distal of soma-to-axon-border; primary: 5.8 μm), but not completely absent (Fig. 5I&L). Strikingly, almost no detectable levels of ANKG were observed at the proximal axon of SH-SY5Y-derived neurons (Fig. 5A,C,G), suggesting the absence of a classical AIS. This is supported by the random-like distribution of ANKG across the axon (Fig. 5M). Further, only low TRIM46 levels were seen across the axon (Fig. 5B,C,H) without significant enrichment at the proximal part (Fig. 5N). This means that SH-SY5Y-derived neurons can efficiently sort endogenous TAU and overexpressed 0N3R-TAU^HA^ without detectable enrichment of ANKG or TRIM46 at the proximal axon. Thus, the process of axonal TAU sorting in neurons may be independent of ANKG and TRIM46 accumulation at the AIS.

## Discussion

Mislocalization of the axonal TAU protein is a hallmark of Alzheimer’s disease and related tauopathies. In this study, we first aimed to establish a suitable neuronal cell model, derived from SH-SY5Y cells, for studying axonal TAU sorting. We then used these SH-SY5Y-derived neurons and mouse primary forebrain neurons i) to evaluate the subcellular sorting of a C-terminus-lacking version of TAU, and ii) to test whether the efficient axonal TAU sorting in SH-SY5Y-derived neurons depends on the enrichment of ANKG and TRIM46 at the proximal axon.

### RA/BDNF-treated SH-SY5Y-derived neurons are suitable for TAU sorting research

In the first step, we evaluated whether SH-SY5Y-derived neurons are a suitable model system for studying axonal TAU sorting. We found that sequential treatment with RA and BDNF generated the highest number of neurons with the most pronounced axonal outgrowth, high expression levels of TAU and MAP2 and efficient axonal sorting of TAU. As seen in former studies^56,79,80^, only the administration of BDNF allowed the use of serum-free media, which explains the pure and thereby stable neuronal cultures. Culture purity, neuronal polarity and TAU sorting efficiency were inferior in SH-SY5Y-derived neurons treated with RA, TPA or TPA. Based on these findings, all further experiments were performed with RA/BDNF-treated SH-SY5Y-derived neurons.

An often-faced problem of rodent primary cultures is the inefficient sorting of transfected and overexpressed TAU constructs^42,50,81^. This effect was also observed in our study, where axonal enrichment of 0N3R-TAU^HA^ was roughly 50 % less than for endogenous TAU in mouse primary neurons. In contrast, the axonal enrichment 0N3R-TAU^HA^ reached endogenous-like levels (~ 94 % of endogenous TAU) in SH-SY5Y-derived neurons. One explanation for this striking difference might be that SH-SY5Y-derived neurons tolerate the overexpression of 0N3R-TAU^HA^ and tdTomato for at least nine days. Successfully transfected primary neurons largely died after more than three days of overexpression (data not shown). The longer expression time may facilitate the axonal targeting of 0N3R-TAU^HA^ as the plasmid DNA may become degraded after several days. This results in a reduction of protein overload and allows more efficient sorting of the exogenous protein. Another cause might be species-specific differences of the TAU sorting machinery, which affects the ability to sort human TAU isoforms in mouse primary neurons^49^. Further, mouse primary neurons might express a different set of isoforms that is more efficiently sorted than 0N3R-TAU. However, 0N3R-TAU was the most efficiently sorted isoform in former studies^41,81^, and 0N3R-TAU is known to be the predominant isoform in early developmental stages^3,82^, probably including div9 neurons derived from E13.5 embryonic mice. For SH-SY5Y-derived neurons, the predominant expression of 0N3R-TAU was reported^62^.

In brief, SH-SY5Y-derived neurons share properties of often-used neuronal models for TAU sorting as rodent primary neurons^40–43^ or iPSC-derived neurons^44–48^. But these cells overcome several of their limitations, as they are human-derived, cultivatable at low cost and without the animal need, and as a neuroblastoma cell line readily accessible for genetic engineering, also within the TAU-encoding *MAPT* gene^68,69^. Moreover, transfected TAU^HA^ is sorted similar to endogenous TAU, which is difficult to achieve in rodent primary cultures^42,50,81^.

### Axonal TAU sorting requires the C-terminal half of TAU

In the next step, we used SH-SY5Y-derived and mouse primary neurons to analyze whether TAU is still efficiently sorted when it lacks the C-terminal half (N-term-TAU^HA^). No axonal enrichment of N-term-TAU^HA^ was seen in both used neuronal cell models. Moreover, N-term-TAU^HA^ tends to accumulate within the soma of transfected neurons. This may hint at the increased accumulation tendency in the absence of the C-terminus or enhanced cellular degradation resulting from protein quality control mechanisms. In the latter case, N-term-TAU^HA^ could be enriched in subcellular compartments, such as lysosomes or stress granules. This somatic localization of N-term-TAU^HA^ is interesting since recent studies claimed a major role for the very N-terminal TAU domain^43^ and the P-rich domain^42^ in mediating axonal TAU sorting. As both domains are present in N-term-TAU^HA^, they are either not involved in TAU sorting or at least not sufficient to maintain the intracellular sorting process without the C-terminus. Notably, the C-terminal repeat domains of TAU are critical for the MT-binding affinity of the entire protein^13,14^. As there is evidence that MTs and MT architecture play a role in the TAU sorting process^34,39,41^, this could explain the necessity of the C-terminus during axonal sorting. However, recent findings question that the C-terminal repeat domains have a strong impact on efficient TAU sorting^42^.

Thus, more effort is required to shed light on the TAU-intrinsic properties that ensure efficient TAU sorting. In this context, SH-SY5Y-derived neurons are valuable to conduct further and more comprehensive studies, e.g. by comparative analyses of differently truncated or modified TAU constructs.

### Axonal TAU sorting is independent of ANKG and TRIM46 enrichment at the proximal axon

The formation of the AIS at the proximal part of the axon is critical for the development and maintenance of neuronal polarity, and for the polarized distribution of proteins^35,37^. There is strong evidence that the AIS and the integrity of its architecture play a major role in regulating the anterograde and retrograde trafficking of TAU^33,34,40,41^.

Therefore, we checked for the formation of an AIS in SH-SY5Y-derived neurons. We first measured the protein levels of Ankyrin G (ANKG), a master organizer of the AIS architecture^35–37^, at the proximal axon of SH-SY5Y-derived and mouse primary neurons. Strikingly, no ANKG was detected within the putative AIS region of SH-SY5Y-derived neurons, in contrast to the mouse primary neurons. The robust axonal sorting of endogenous and transfected TAU does apparently not require ANKG enrichment at the proximal axon or detectable ANKG expression at all although ANKG is responsible for the recruitment of major AIS structural components^35^. The tracing of MAP2 signal revealed invasion of MAP2 into the proximal axon. A lack of MAP2 polarity is reported for ANKG-deficient cultured neurons^38,83^. However, the low MAP2 levels more distal than 50 μm clearly prove a still existing, albeit reduced, ANKG-independent somatic MAP2 retention in these cells (see Figs. 2&5).

Recent studies revealed the tripartite motif-containing protein 46 (TRIM46), an axonal MT-organizing protein^39,84^, as a major player for the initial development of neuronal polarity^39^. TRIM46 accumulates at the proximal axon prior to ANKG and forms highly polarized MT fascicles^38,39^. TRIM46-deficiency leads to TAU missorting in cultured primary neurons^77^. Therefore, we investigated the TRIM46 levels at the proximal axon of SH-SY5Y-derived neurons. While the mouse primary neurons showed a clearly defined TRIM46 accumulation at the AIS, there were only small amounts of TRIM46 protein along the axon without a significant enrichment at the putative AIS region.

The absence of a classical AIS, indicated by the lack of ANKG and TRIM46 enrichment, together with the predominant 0N3R isoform expression^62^ might hint at the incomplete maturity of SH-SY5Y-derived neurons^35^. However, SH-SY5Y-derived neurons express several neuronal maturation marker proteins^56–62^, and there are neuron subtypes known to lack ANKG-dependent AIS formation as well^85^. The spatial separation of endogenous TAU and MAP2 in SH-SY5Y-derived neurons is slightly less efficient compared to mouse primary neurons, which could be a direct consequence of the ANKG and TRIM46 absence, as both were shown to promote somatic retention of MAP2^83^ and axonal sorting of TAU directly^39,77^.

As ANKG and TRIM46 are absent and thus do not mediate axonal sorting of TAU in SH-SY5Y-derived neurons, the question arises, which remaining factors mediate the robust axonal targeting of TAU seen in these neurons. The end-binding protein 3 (EB3) was recently shown to be critical for the AIS integrity^86,87^. EB3 accumulates and binds along the MT network at the AIS^87,88^, and links it to ANKG. This EB3-ANKG interplay is necessary for the proper assembly and maintenance of the AIS by regulating the MT architecture^86,87^. As EB3 also binds to the plus end of growing axons, it could directly promote anterograde trafficking of TAU and other axonal proteins via a piggyback-like mechanism^89,90^. Interestingly, TAU and EB3 were recently shown to co-localize at the AIS of iPSC-derived cortical neurons^48^. A central role in AIS assembly and maintenance was also shown for the MT-binding protein MTCL-1^91^.

The relevance of TRIM46 and other MT-binding proteins in maintaining neuronal polarity^39,86,87,91^ strongly suggest that the organization and regulation of the MT network at the AIS is one key factor for its structural and functional integrity, which includes the correct intracellular sorting of MAP2, TAU and other compartment-specific proteins^38^. However, our results question the necessity of TRIM46 for the development of neuronal polarity and polarized Tau distribution. Interestingly, TAU and EB3 co-localize at the AIS of iPSC-derived cortical neurons^48^. It is still unclear whether this co-localization is relevant for TAU trafficking or due to the MT affinity of both proteins. Taken together, the role of MT-organizing proteins in the context of AIS integrity and axonal TAU sorting remains inconclusive and needs further elaboration, with SH-SY5Y-derived neurons as a potential tool.

### Conclusion

In this study, we demonstrate that SH-SY5Y-derived human neurons, generated with RA/BDNF treatment, are a suitable tool for investigating TAU sorting mechanisms in human cells. These neuronal cells show pronounced neuronal polarity, neuronal marker expression, efficient sorting of endogenous and, importantly, of transfected TAU and TAU constructs similar to mouse primary neurons. By using SH-SY5Y-derived neurons, we show that the N-terminal half of TAU is not sufficient for successful axonal sorting. Further, axonal TAU targeting is independent of classical AIS formation, as ANKG and TRIM46 enrichment at the proximal axon is absent in SH-SY5Y derived neurons, but TAU is still efficiently sorted into the axon. Taken together, our data show i) that the C-terminal half of TAU is required for efficient axonal sorting, and ii) that subcellular sorting of TAU might be largely independent of classical AIS formation, which questions the claimed role of ANKG, TRIM46 and other MT-organizing proteins in developing and maintaining neuronal polarity.

## Acknowledgments

We thank Prof. Dr. Rudolf Wiesner (Institute for Veg. Physiology II, University Hospital Cologne) for providing us the SH-SY5Y neuroblastoma cell line, and Dr. Magdalena Bogus (Institute for Forensic Medicine) for cell authentication. Animals were obtained from the CMMC animal facility and the CECAD in vivo research facility (both Cologne, Germany). This work was funded by the Else-Kröner-Fresenius Stiftung, Cologne Fortune, and supported by a doctoral fellowship of the Studienstiftung des deutschen Volkes. The authors declare that they have no competing interests.

## Author contributions

MB: study design, experimental conduct, data acquisition, analysis, and interpretation, manuscript drafting. SB: manuscript proofreading, assistance in methodology development. JK: experimental conduct, assistance in data acquisition and analysis. FL: assistance in experimental conduct (molecular biology). HZ: project funding, study design, data interpretation, manuscript drafting.

## Notes

### Competing Interest Statement

The authors have declared no competing interest.

### Summary of Updates

The plot profiles of MAP2 and the AIS-located protein TRIM46 were added to Figure 5. The distribution of ANKG and TRIM46 along the proximal axon were quantified.

