## Supplemental Material for "Axonal TAU sorting requires the C-terminus of TAU but is independent of ANKG and TRIM46 enrichment at the AIS"

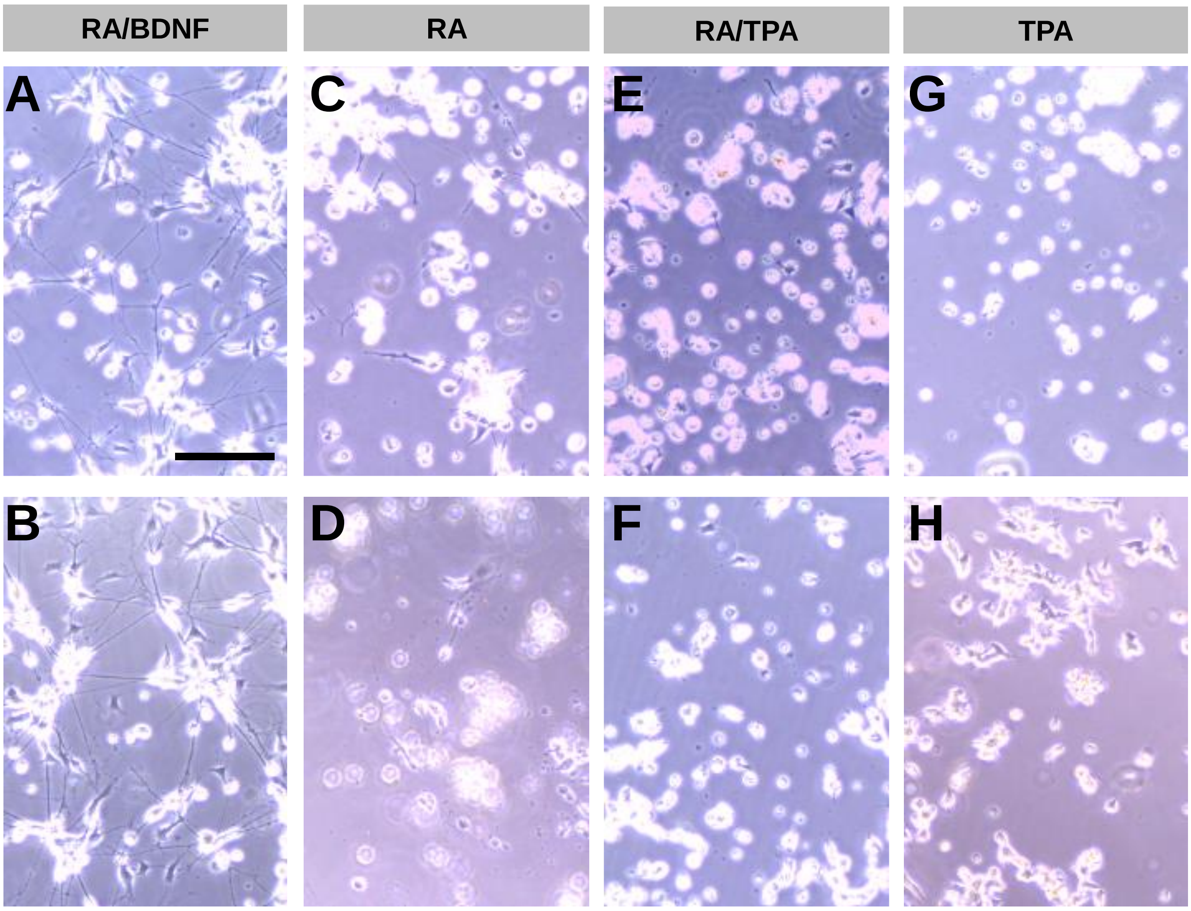


**Supplemental Figure 1: Alternative differentiation protocols for RA, RA/TPA and TPA**. **A&B:** Representative culture images of RA/BDNF-treated SH-SY5Y-derived neurons (d14) after transfection with tdTomato at d5 (see methods for details). Note the viable cultures without increased cell death compared to untransfected cultures (see Fig. 1C). Scale bar: 50 µm. **C-H:** Representative culture images of RA- (C&D), RA/TPA- (E&F) and TPA- (G&H) treated SH-SY5Y-derived neurons. Cultures were differentiated by using two alternative long-term protocols until d14, namely variant A with identical seeding density and stepwise serum reduction (C,E,G), and variant B with increased seeding density and stepwise serum reduction (D,F,H) (see methods for details). The cultivation without serum should prevent excessive proliferation of undifferentiated SH-SY5Y cells. Note the sparse cultures and the high number of detached cells compared to RA/BDNF-treated cultures after transfection (see Fig. 1C, S1A&B).

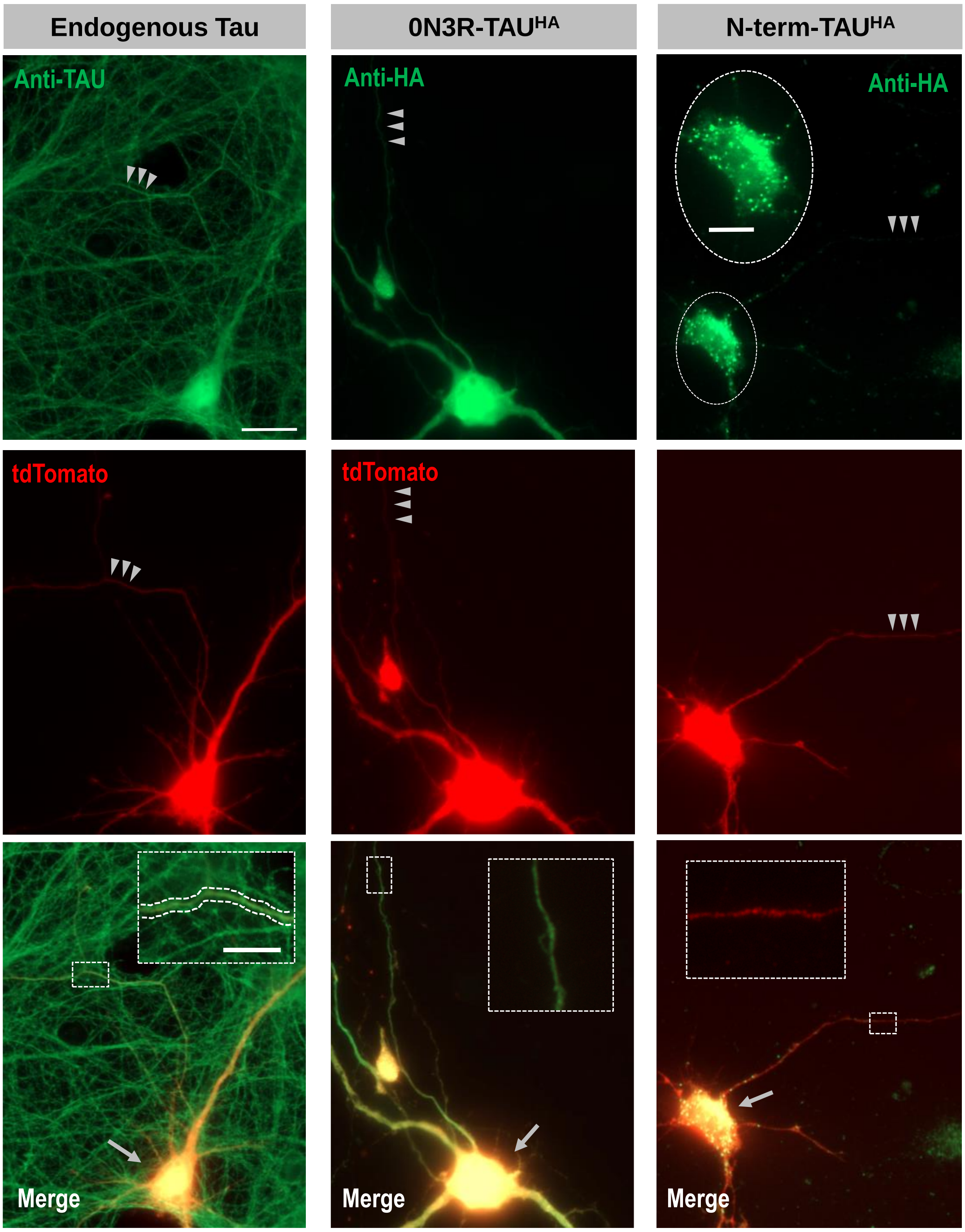


**Supplemental Figure 2: Axonal enrichment of endogenous TAU and transfected TAU$^{HA}$ constructs in mouse primary neurons.** Representative immunofluorescent images of mouse primary neurons transfected with tdTomato (left column), or tdTomato and 0N3R-TAU$^{HA}$ (middle column) or N-term-TAU$^{HA}$ (right column). Endogenous TAU (A, green) or TAU$^{HA}$ constructs (B, C, green) and co-transfected tdTomato as volume marker (red) are shown in single channels and merged. Cells were transfected at div6 and fixed at div9 (see methods for details). Note the strong axonal sorting of endogenous TAU compared to tdTomato (arrowheads), which leads to yellowish somatic staining (arrows) and mainly green axonal staining in merged images. The axonal sorting of 0N3R-TAU$^{HA}$ is less pronounced (see Fig. 4C for quantification). Axons of cells with N-term-TAU$^{HA}$ appear red, indicating weak axonal sorting of N-term-TAU$^{HA}$. Axonal sections (small dashed boxes) are shown with higher magnification (large dashed boxes, dashed lines trace the axon in the left merge image). HA-positive puncta (small dashed circle, magnified in large dashed circle) appear in many N-term-TAU$^{HA}$ transfected cells but not in cells expressing 0N3R-TAU$^{HA}$. Scale bar: 20 µm, scale bar within dashed box: 3 µm.

**Supplemental Sequence 1:** Sequence of the CMV-driven expression vector with the 0N3R-TAU$^{HA}$ insert (pHA-0N3R-TAU_HA)

TAGTTATTAATAGTAATCAATTACGGGGTCATTAGTTCATAGCCCATATATGGAG-
TTCCGCGTTACATAACTTACGGTAAATGGCCCGCCTGGCTGACCGCCCAACGACCCCCGCCCAT
TGACGTCAATAATGACGTATGTTCCCATAGTAACGCCAATAGGGACTTTCCATTGAC-
GTCAATGGGTGGAGTATTTACGGTAAACTGCCCACTTGGCAGTACATCAAGTGTATCATATGCCA
AGTACGCCCCCTATTGACGTCAATGACGGTAAATGGCCCGCCTGGCATTATGCCCAG-
TACATGACCTTATGGGACTTTCCTACTTGGCAGTACATCTACGTATTAGTCATCGCTATTACCATG
GTGATGCGGTTTTGGCAGTACATCAATGGGCGTGGATAGCGGTTTGACTCACGGGGAT-
TTCCAAGTCTCCACCCCATTGACGTCAATGGGAGTTTGTTTTGGCACCAAAATCAACGGGACTTT
CCAAAATGTCGTAACAACTCCGCCCCATTGACGCAAATGGGCGGTAGGCGTGTACGGTGG-
GAGGTCTATATAAGCAGAGCTGGTTTAGTGAACCGTCAGATCCGCTAGCGCTACCGGTGCCACC
ATGTACCCATACGATGTTCCAGATTACGCTGAGCCCCGCCAGGAGTTCGAAGTGATGGAA-
GATCACGCTGGGACGTACGGGTTGGGGGACAGGAAAGATCAGGGGGGCTACACCATGCACCAA
GACCAAGAGGGTGACACGGACGCTGGCCTGAAAGAATCTCCCCTGCAGACCCCCACTGAG-
GACGGATCTGAGGAACCGGGCTCTGAAACCTCTGATGCTAAGAGCACTCCAACAGCGGAAGATG
TGACAGCACCCTTAGTGGATGAGGGAGCTCCCGGCAAGCAGGCTGCCGCGCAGCCCCACAC-
GGAGATCCCAGAAGGAACCACAGCTGAAGAAGCAGGCATTGGAGACACCCCCAGCCTGGAAGA
CGAAGCTGCTGGTCACGTGACCCAAGCTCGCATGGTCAGTAAAAGCAAAGACGGGACTGGAA-
GCGATGACAAAAAAGCCAAGGGGGCTGATGGTAAAACGAAGATCGCCACACCGCGGGGAGCAG
CCCCTCCAGGCCAGAAGGGCCAGGCCAACGCCACCAGGATTCCAG-
CAAAAACCCCGCCCGCTCCAAAGACACCACCCAGCTCTGGTGAACCTCCAAAATCAGGGGATCG
CAGCGGCTACAGCAGCCCCGGCTCCCCAGGCACTCCCGGCAGCCGCTCCCG-
CACCCCGTCCCTTCCAACCCCACCCACCCGGGAGCCCAAGAAGGTGGCAGTGGTCCGTACTCC
ACCCAAGTCGCCGTCTTCCGCCAAGAGCCGCCTGCAGACAGCCCCCGTGCCCATGCCAGAC-
CTGAAGAATGTCAAGTCCAAGATCGGCTCCACTGAGAACCTGAAGCACCAGCCGGGAGGCGGG
AAGGTGCAGATAATTAATAAGAAGCTGGATCTTAGCAACGTCCAG-
TCCAAGTGTGGCTCAAAGGATAATATCAAACACGTCCCGGGAGGCGGCAGTGTGCAAATAGTCT
ACAAACCAGTTGACCTGAGCAAGGTGACCTCCAAGTGTGGCTCATT-
AGGCAACATCCATCATAAACCAGGAGGTGGCCAGGTGGAAGTAAAATCTGAGAAGCTTGACTTCA
AGGACAGAGTCCAGTCGAAGATTGGGTCCCTGGACAATATCACCCACGTCCCTGGCGGAG-
GAAATAAAAAGATTGAAACCCACAAGCTGACCTTCCGCGAGAACGCCAAAGCCAAGACAGACCA
CGGGGCGGAGATCGTGTACAAGTCGCCAGTGGTGTCTGGGGACACGTCTCCACGG-
CATCTCAGCAATGTCTCCTCCACCGGCAGCATCGACATGGTAGACTCGCCCCAGCTCGCCACGC
TAGCTGACGAGGTGTCTGCCTCCCTGGCCAAGCAGGGTTTGTGAGGATCCACCGGATCTAGA-
TAACTGATCATAATCAGCCATACCACATTTGTAGAGGTTTTACTTGCTTTAAAAAACCTCCCACACC
TCCCCCTGAACCTGAAACATAAAATGAATGCAATTGTTGTTGTTAACTT-
GTTTATTGCAGCTTATAATGGTTACAAATAAAGCAATAGCATCACAAATTTCACAAATAAAGCATTT
TTTTCACTGCATTCTAGTTGTGGTTTGTCCAAACTCATCAATGTATCTTAACGCGTAAATT-
GTAAGCGTTAATATTTTGTTAAAATTCGCGTTAAATTTTTGTTAAATCAGCTCATTTTTTAACCAATA
GGCCGAAATCGGCAAAATCCCTTATAAATCAAAAGAATAGACCGAGATAGGGTTGAGTGTT-
GTTCCAGTTTGGAACAAGAGTCCACTATTAAAGAACGTGGACTCCAACGTCAAAGGGCGAAAAAC
CGTCTATCAGGGCGATGGCCCACTACGTGAACCATCACCCTAATCAAGTTTTTT-
GGGGTCGAGGTGCCGTAAAGCACTAAATCGGAACCCTAAAGGGAGCCCCCGATTTAGAGCTTGA
CGGGGAAAGCCGGCGAACGTGGCGAGAAAGGAAGGGAAGAAA-
GCGAAAGGAGCGGGCGCTAGGGCGCTGGCAAGTGTAGCGGTCACGCTGCGCGTAACCACCACA
CCCGCCGCGCTTAATGCGCCGCTACAGGGCGCGTCAGGTGGCACTTTTCGGG-
GAAATGTGCGCGGAACCCCTATTTGTTTATTTTTCTAAATACATTCAAATATGTATCCGCTCATGAG
ACAATAACCCTGATAAATGCTTCAATAATATTGAAAAAGGAAGAGTCCTGAGGCGGAAA-
GAACCAGCTGTGGAATGTGTGTCAGTTAGGGTGTGGAAAGTCCCCAGGCTCCCCAGCAGGCAGA
AGTATGCAAAGCATGCATCTCAATTAGTCAGCAACCAGGTGTG-
GAAAGTCCCCAGGCTCCCCAGCAGGCAGAAGTATGCAAAGCATGCATCTCAATTAGTCAGCAAC
CATAGTCCCGCCCCTAACTCCGCCCATCCCGCCCCTAACTCCGCCCAG-
TTCCGCCCATTCTCCGCCCCATGGCTGACTAATTTTTTTTATTTATGCAGAGGCCGAGGCCGCCT
CGGCCTCTGAGCTATTCCAGAAGTAGTGAGGAGGCTTTTTTGGAGGCCTAGGCTTTTGCAAA-
GATCGATCAAGAGACAGGATGAGGATCGTTTCGCATGATTGAACAAGATGGATTGCACGCAGGTT

CTCCGGCCGCTTGGGTGGAGAGGCTATTCGGCTATGACTGGGCACAACAGA-
CAATCGGCTGCTCTGATGCCGCCGTGTTCCGGCTGTCAGCGCAGGGGCGCCCGGTTCTTTTTGT
CAAGACCGACCTGTCCGGTGCCCTGAATGAACTGCAAGACGAGGCAGCGCGGC-
TATCGTGGCTGGCCACGACGGGCGTTCCTTGCGCAGCTGTGCTCGACGTTGTCACTGAAGCGG
GAAGGGACTGGCTGCTATTGGGCGAAGTGCCGGGGCAGGATCTCCTGTCATCTCACCTT-
GCTCCTGCCGAGAAAGTATCCATCATGGCTGATGCAATGCGGCGGCTGCATACGCTTGATCCGG
CTACCTGCCCATTCGACCACCAAGCGAAACATCGCATCGAGCGAGCACGTACTCGGATGGAA-
GCCGGTCTTGTCGATCAGGATGATCTGGACGAAGAGCATCAGGGGCTCGCGCCAGCCGAACTG
TTCGCCAGGCTCAAGGCGAGCATGCCCGACGGCGAG-
GATCTCGTCGTGACCCATGGCGATGCCTGCTTGCCGAATATCATGGTGGAAAATGGCCGCTTTTC
TGGATTCATCGACTGTGGCCGGCTGGGTGTGGCGGACCGCTATCAGGACATAGCGTTGGC-
TACCCGTGATATTGCTGAAGAGCTTGGCGGCGAATGGGCTGACCGCTTCCTCGTGCTTTACGGT
ATCGCCGCTCCCGATTCGCAGCGCATCGCCTTCTATCGCCTTCTTGACGAG-
TTCTTCTGAGCGGGACTCTGGGGTTCGAAATGACCGACCAAGCGACGCCCAACCTGCCATCACG
AGATTTCGATTCCACCGCCGCCTTCTATGAAAGGTTGGGCTTCGGAATCGTTTTCCGGGAC-
GCCGGCTGGATGATCCTCCAGCGCGGGGATCTCATGCTGGAGTTCTTCGCCCACCCTAGGGGG
AGGCTAACTGAAACACGGAAGGAGACAATACCGGAAGGAACCCGCGCTATGACGGCAA-
TAAAAAGACAGAATAAAACGCACGGTGTTGGGTCGTTTGTTCATAAACGCGGGGTTCGGTCCCAG
GGCTGGCACTCTGTCGATACCCCACCGAGACCCCATTGGGGCCAATAC-
GCCCGCGTTTCTTCCTTTTCCCCACCCCACCCCCCAAGTTCGGGTGAAGGCCCAGGGCTCGCAG
CCAACGTCGGGGCGGCAGGCCCTGCCATAGCCTCAGGTTACTCATATATACTTTAGATTGAT-
TTAAAACTTCATTTTTAATTTAAAAGGATCTAGGTGAAGATCCTTTTTGATAATCTCATGACCAAAAT
CCCTTAACGTGAGTTTTCGTTCCACTGAGCGTCAGACCCCGTAGAAAA-
GATCAAAGGATCTTCTTGAGATCCTTTTTTTCTGCGCGTAATCTGCTGCTTGCAAACAAAAAAACC
ACCGCTACCAGCGGTGGTTTGTTTGCCGGATCAAGAGCTAC-
CAACTCTTTTTCCGAAGGTAACTGGCTTCAGCAGAGCGCAGATACCAAATACTGTCCTTCTAGTGT
AGCCGTAGTTAGGCCACCACTTCAAGAACTCTGTAGCACCGCCTACAT-
ACCTCGCTCTGCTAATCCTGTTACCAGTGGCTGCTGCCAGTGGCGATAAGTCGTGTCTTACCGG
GTTGGACTCAAGACGATAGTTACCGGATAAGGCGCAGCGGTCGGGCTGAAC-
GGGGGGTTCGTGCACACAGCCCAGCTTGGAGCGAACGACCTACACCGAACTGAGATACCTACAG
CGTGAGCTATGAGAAAGCGCCACGCTTCCCGAAGGGAGAAAGGCGGACAGGTATCCGGTAA-
GCGGCAGGGTCGGAACAGGAGAGCGCACGAGGGAGCTTCCAGGGGGAAACGCCTGGTATCTTT
ATAGTCCTGTCGGGTTTCGCCACCTCTGACTTGAGCGTCGATTTTT-
GTGATGCTCGTCAGGGGGGCGGAGCCTATGGAAAAACGCCAGCAACGCGGCCTTTTTACGGTTC
CTGGCCTTTTGCTGGCCTTTTGCTCACATGTTCTTTCCTGCGTTATCCCCTGATTCTGTGGA-
TAACCGTATTACCGCCATGCAT
